# DNA-PKcs phosphorylation at the T2609 cluster alters the repair pathway choice during immunoglobulin class switch recombination

**DOI:** 10.1101/2020.04.23.057877

**Authors:** Jennifer L. Crowe, Xiaobin S. Wang, Zhengping Shao, Brian J. Lee, Verna Estes, Shan Zha

**Affiliations:** Institute for Cancer Genetics, Vagelos College of Physicians and Surgeons, Columbia University, New York City, NY 10032; Graduate program of Pathobiology and Molecular Medicine, Vagelos College of Physicians and Surgeons, Columbia University, New York City, NY 10032; Division of Pediatric Oncology, Hematology and Stem Cell Transplantation, Department of Pediatrics, Vagelos College of Physicians & Surgeons, Columbia University, New York City, NY 10032

## Abstract

The DNA-dependent protein kinase (DNA-PK), composed of the KU heterodimer and the large catalytic subunit (DNA-PKcs), is a classical non-homologous end-joining (cNHEJ) factor. Naïve B cells undergo class switch recombination (CSR) to generate antibodies with different isotypes by joining two DNA double-strand breaks at different switching regions via the cNHEJ pathway. DNA-PK and the cNHEJ pathway play important roles in the DNA repair phase of CSR. To initiate cNHEJ, KU binds to DNA ends, and recruits and activates DNA-PK. DNA-PKcs is the best-characterized substrate of DNA-PK, which phosphorylates DNA-PKcs at both the S2056 and T2609 clusters. Loss of T2609 cluster phosphorylation increases radiation sensitivity, suggesting a role of T2609 phosphorylation in DNA repair. Using the *DNA-PKcs^5A^* mouse model carrying an alanine substitution at the T2609 cluster, here we show that loss of T2609 phosphorylation of DNA-PKcs does not affect the CSR efficiency. Yet, the CSR junctions recovered from *DNA-PKcs^5A/5A^* B cells reveal increased chromosomal translocation, excess end-resection, and preferential usage of micro-homology – all signs of the alternative end-joining pathway. Thus, these results uncover a role of DNA-PKcs T2609 phosphorylation in promoting cNHEJ repair pathway choice during CSR.

**Key points:** Loss of T2069 cluster phosphorylation of DNA-PKcs promotes Alt-EJ-mediated CSR.

## Introduction

DNA-dependent protein kinase (DNA-PK), composed of the KU70 and KU86 (Ku80 in mouse) heterodimer (KU) and a large catalytic subunit (DNA-PKcs), is a member of the classical non-homologous end-joining (cNHEJ) pathway. As a major DNA double-strand break (DSB) repair pathway in mammalian cells, cNHEJ entails both end-processing and end-ligation. KU initiates cNHEJ by binding to the double-stranded DNA (dsDNA) ends, which in turn recruits and activates DNA-PKcs that, among other functions, activates Artemis endonuclease for end-processing (1, 2). KU also interacts with and stabilizes the LIG4/XRCC4/XLF complex for end-ligation. In animal models, loss of cNHEJ mediated end-ligation (*e.g., Lig4^-/-^*) leads to severe neuronal apoptosis, which causes embryonic lethality in the case of *Xrcc4^-/-^* and *Lig4^-/-^* mice (3). In contrast, loss of DNA-PKcs or Artemis increases radiation sensitivity but does not cause measurable neuronal apoptosis, consistent with their relatively limited roles in direct end-ligation, if any (4–6). Several new cNHEJ factors (*e.g*., PAXX, and CYREN/MRI) are characterized based on their interaction with KU. Loss of PAXX or CYREN/MRI increases radiation sensitivity, but only abrogates end-ligation if XLF, a non-essential cNHEJ factor, is also lost simultaneously (7–11).

In addition to general DSB repair, developing lymphocytes undergo programmed DSB break and repair events to assemble and modify the immunoglobulin heavy chain (IgH) genes. Specifically, V(D)J recombination, which occurs in progenitor lymphocytes, assembles the variable region exon from individual V, D, and J gene segments exclusively via cNHEJ. DNA-PKcs and Artemis are required for opening the hairpin at the coding ends during V(D)J recombination. Correspondingly, *DNA-PKcs^-/-^* (null) or *Artemis^-/-^* mice are born of normal size at the expected ratio but develop severe combined immunodeficiency (SCID) (4–6). Patients with hypomorphic mutations in DNA-PKcs or Artemis also develop SCID (12, 13). Mature B cells undergo CSR, which replace the initially expressed IgM constant region with another downstream constant region encoding a different isotype, to generate antibodies with different effector functions. The cNHEJ pathway plays an important role in CSR. But in cells lacking a core cNHEJ factor (*e.g*., Lig4 or Xrcc4), up to 25-50% of CSR can be achieved by the alternative end-joining (Alt-EJ) pathway that preferentially uses microhomology (MH) at the junctions. Consistent with DNA-PKcs and Artemis being dispensable for direct end-ligation, *DNA-PKcs^-/-^* B cells only have moderate defects in CSR (14, 15). Nevertheless, in recent high-throughput sequence analyses, we found that CSR junctions recovered from *DNA-PKcs^-/-^* B cells contained increased chromosomal translocations and extensive end-resection, and preferentially used MHs (16), suggesting that the seemingly robust CSR achieved in *DNA-PKcs^-/-^* cells is primarily mediated by the Alt-EJ pathway like those in *Lig4^-/-^* or *Xrcc4^-/-^* B cells (17, 18). Accordingly, human patients with spontaneous mutations in DNA-PKcs show severe defects in both V(D)J recombination and CSR and increased MH in the residual CSR junctions(12, 19).

Active DNA-PK phosphorylates many substrates, among which, DNA-PKcs is the best-characterized (20). In a previous study, we showed that mice carrying a knock-in kinase-dead mutation of DNA-PKcs (*DNA-PKcs^KD^*) die *in utero* with severe neuronal apoptosis, like *Lig4^-/-^* or *Xrcc4^-/-^* mice. KU deletion rescued the embryonic development of *DNA-PKcs^KD/KD^* mice, suggesting that DNA-PKcs has an end-capping function at the DNA ends that is regulated by its kinase activity. Correspondingly, *DNA-PKcs^KD/KD^* mature B cells display severe CSR defects like *Lig4^-/-^* B cells (16) and the residual CSR junctions from *DNA-PKcs^KD/KD^* B cells are all mediated by the Alt-EJ pathway

Three DNA damage-induced phosphorylation clusters on DNA-PKcs have been characterized: the S2056 cluster, the T2609 cluster, and the S3590 cluster (Fig. 1A)(21–24). Among them, the T2609 cluster can also be cross phosphorylated by ATM upon ionizing radiation (25, 26) and by ATR upon UV radiation (27). Human DNA-PKcs with alanine substitutions at the S2056 cluster and/or the T2609 cluster, cannot restore radiation resistance in DNA-PKcs-deficient Chinese hamster ovary (CHO) cells (22, 28–31), suggesting a role of DNA-PKcs phosphorylation in DSB repair. The mouse model carrying alanine substitutions in the S2056 cluster (*DNA-PKcs^PQR/PQR^*) has no defects in chromosomal V(D)J Recombination or CSR, despite moderate radiation sensitivity (32). Mouse models with alanine substitutions of three or all five threonines at the T2609 cluster (*DNA-PKcs^3A/3A^ or DNA-PKcs^5A/5A^*) developed neonatal bone marrow failure that abolishes both myeloid and lymphoid progenitors (33–35). *Tp53*-deficiency restores lymphocyte development and peripheral mature lymphocyte counts in *DNA-PKcs^3A/3A^* and *DNA-PKcs^5A/5A^* mice and v-abl kinase transformed *DNA-PKcs^3A/3A^* and *DNA-PKcs^5A/5A^* B cells undergo robust chromosomal V(D)J recombination in culture (35, 36), suggesting that T2609 phosphorylation is not required for V(D)J recombination.

**Figure 1:**
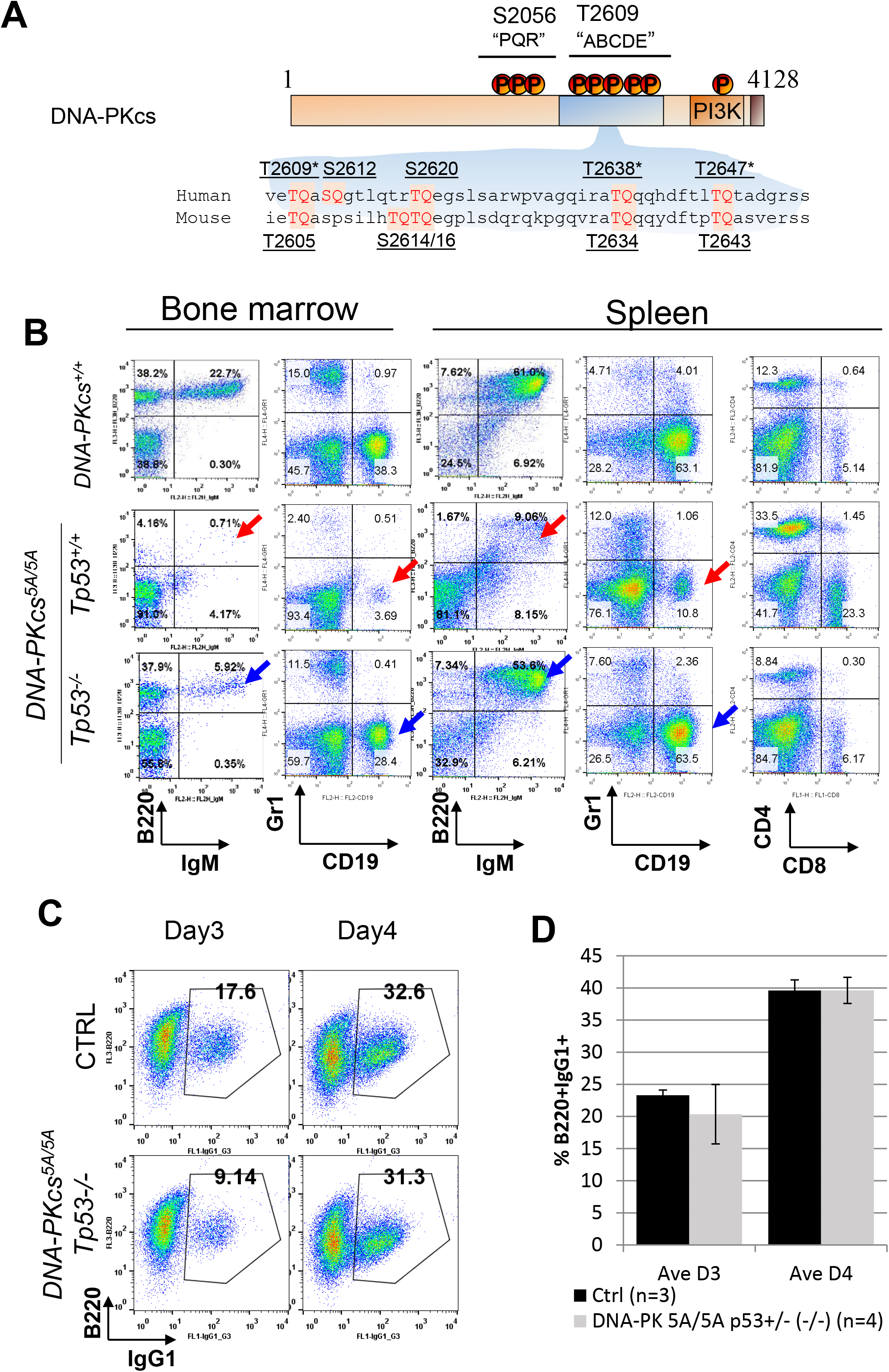
Class switch recombination is transiently delayed in *DNA-PKcs^5A/5A^* cells. (A) Diagram of the DNA-PKcs protein and the organization of the T2609 cluster on human DNA-PKcs and the corresponding T2605 cluster on mouse DNA-PKcs. The conserved T2609, T2636 and T2647 are marked with asterisks. The *DNA-PKcs^5A^* allele contains alanine substitutions of all 5 threonines, while the previously published DNA-PKcs 3A allele contains alanine substitions at three conserved threonines. (B) Representative flow cytometry analyses of B and T cell development in *DNA-PKcs^5A/5A^* and *DNA-PKcs^5A/5A^Tp53^-/-^* mice, indicating *Tp53*-deficiency restored peripheral mature B cells (IgM+B220+) in *DNA-PKcs^5A/5A^* mice. (C) Representative flow cytometry analyses of class switch recombination to IgG1 at day 3 and day 4 after cytokine activation. (D) Quantification of IgG1 switching. The bars mark the standard erros. There is no significant difference between *DNA-PKcs^5A/5A^* and *DNA-PKcs^+/+^* B cells.

Here we analyzed the impact of DNA-PKcs T2609 phosphorylation on CSR in mature B cells isolated from *DNA-PKcs^5A/5A^* mice carrying *Tp53*-deficiency (heterozygous or homozygous). While CSR efficiency is not affected by T2609A substitution, high throughput sequence analyses reveal that the CSR junctions from *DNA-PKcs^5A/5A^* B cells contain increased chromosomal translocations, evidence of excessive end-resection, and increased MHs – all signatures of the Alt-EJ pathway. Therefore, our findings uncover a role of DNA-PKcs T2609 phosphorylation in repair pathway choice during CSR.

## Materials and Methods

### Generation and characterization of the mouse models

*DNA-PKcs^-/-^*, *p53^-/-^* and the pre-arranged IgH heavy and light chain knock-in alleles (HL^k/k^) that bypass the need for DNA-PKcs in early B cell development, have been described previously (4, 37–39). The *DNA-PKcs^5A^* allele and the generation of the *DNA-PKcs^5A/5A^* mouse model were recently published (35). The *DNA-PKcs^5A^* mutation converts all 5 threonines in the T2605 cluster (T2609 in human) encoded by exon 58 of murine *DNA-PKcs* to alanines (Fig.1A). Briefly, DNA sequences (5’ and 3’ arms) from the genomic DNA-PKcs (*Prkdc*) locus were isolated via PCR and cloned into the pGEMT vector. We synthesized a ~500bp fragment containing all of the mutations (Genewiz), inserted it into an extended 3’arm (4.4kb) in pGEMT, and confirmed by sequencing. The 5’ arm and the extended 3’arm with the mutations were then inserted into the pEMCneo vector that carries a Neo-Resistance (NeoR) gene flanked by a pair of FRT sites. The Sma1 linearized plasmids were electroporated into murine ES cells (129Sv background) and NeoR^+^ clones were isolated and screened by Southern blotting and Sanger sequencing (Fig.1A) (35). Upon injection and germline transmission, the resultant DNA-PKcs^+/5AN^ mice (N for NeoR containing) were crossed with constitutively FLIpase expressing *Rosa26a^FLIP/FLIP^* mice (Jax stock number 003946, also in 129Sv background) to remove the NeoR cassette and allow the expression of the DNA-PKcs-5A allele. All animal work was conducted in a specific pathogen-free facility and all the procedures were approved by the Institutional Animal Care and Use Committee (IACUC) at Columbia University Medical Center.

### Lymphocyte development analyses

Single-cell suspensions were prepared from the thymus, bone marrow, and spleen of young adult (6-8 weeks) mice. Splenocytes were treated with red blood cell lysis buffer (Lonza ACK Lysis Buffer) for 1-2 minutes at room temperature. Approximately 1 × 10^5^ cells were stained using fluorescence-conjugated antibodies and analyzed by flow cytometry (11, 40). The following antibody cocktails were used for B cell (FITC anti-mouse CD43, Biolegend, 553270; PE Goat anti-mouse IgM, Southern Biotech, 1020-09; PE-Cyanine5 anti-Hu/Mo CD45R (B220), eBioScience, 15-0452-83; and APC anti-mouse TER119, Biolegend, 116212) and T cell (PE rat anti-mouse CD4, Biolegend, 557308; FITC anti-mouse CD8a, Biolegend, 100706; PE/Cy5 anti-mouse CD3e, eBioscience, 15-0031-83; and APC anti-mouse TCRβ, BD Pharmingen, 553174) analyses. The dead cells and debris were excluded based on their high side scatter and low forward scatter. For bone marrow and spleen with significant and sometimes variable erythrocytes remaining after red blood cell lysis, we used the Ter119 (an erythrocyte marker) antibody to remove the red blood cells before the analyses of B, T, and myeloid cell-specific markers as in Figure 1.

### Class switch recombination analyses

For the CSR assay, CD43^-^ splenocytes were isolated with magnetic beads (MACS, Miltenyi Biotec) and cultured (5-10 x 10^5^ cells ml^-1^) in RPMI (GIBCO) supplemented with 15% FBS (Hyclone) and 2 μg ml^-1^ anti-CD40 (BD Bioscience) plus 20 ng ml^-1^ of IL-4 (R&D). Cells were maintained daily at 1 x 10^6^ cells ml^-1^ and collected for flow cytometry after staining for IgG1 and B220 (FITC anti-IgG1, BD Pharmingen, and PE Cy5 anti-B220, eBioscience). Flow cytometry data were collected on a FACSCalibur flow cytometer (BD Bioscience) and processed using the FlowJo V10 software package. For the CTV staining, purified B cells were incubated at 6 × 10^6^ cells ml^-1^ in 1 mL of PBS with 1 uL of 5 mM Cell Trace Violet (CTV) dye (Thermo Scientific). The CTV-stained B cells were washed and activated as detailed above and subjected to flow cytometry analyses 4 days after activation.

### High throughput genomic translocation sequencing (HTGTS) of CSR junctions

HTGTS was performed as described (16, 41). Briefly, genomic DNA was collected from activated B cells 4 days after stimulation, sonicated (Diagenode Bioruptor), and amplified with an Sμ specific biotinylated primer (5’/5BiosG/CAGACCTGGGAATGTATGGT3’) and nested primer (5’CACACAAAGACTCTGGACCTC3’). AflII was used to remove germline (non-rearranged) sequences. Since all mice were of pure 129 background, the IgH switch region (from JH4 to the last Cα exon, chr12: 114,494,415-114,666,816) of the C57/BL6 based mm9 genome was replaced with the corresponding region of the AJ851868.3 129 IgH sequence (1415966-1592715) to generate the mm9sr (switch region replacement) genome and the sequence analyses were performed as detailed previously (41, 42). The best-path searching algorithm (related to YAHA (43)) was used to identify optimal sequence alignments from Bowtie2-reported top alignments (alignment score > 50). The reads were filtered to exclude mispriming, germline (unmodified), sequential joints, and duplications. To plot all of the S-region junctions, including those within the repeats and thus have low mappability but unequivocally mapped to an individual switch region, we combined the ones filtered by a mappability filter but unequivocally mapped to S regions with ‘good’ reads passing both the mappability filters (both de-duplicated) (41). MHs are defined as regions of 100% homology between the bait and prey-break sites. Insertions are defined as regions containing nucleotides that map to neither the bait nor prey break site. Blunt junctions are considered to have no MHs or insertions. We calculated the AGCT or RGYW motif frequency in each switch region using the IgH region sequence from the 129/Sv strain (accession number AJ851868.3). The mutation rate was calculated by a customized Excel integrated VBA script. Specifically, we analyzed the mutations in the bait portion by comparing the actual bait sequence of each IgH junction with the germline bait sequence from the 129/Sv strain (accession number AJ851868.3). Only true mismatches (no insertion or deletion) were counted as mutations. The HTGTS data reported in this paper have been deposited in the Gene Expression Omnibus (GEO) database, https://www.ncbi.nlm.nih.gov/geo/ (accession no. GSE117628 and others).

## Results

### Loss of the Tp53 tumor suppressor gene restores peripheral B cells in *DNA-PKcs^5A/5A^* mice

The human DNA-PKcs T2609 phosphorylation cluster contains five TQ sites, among which, T2609, T2638 and T2647 (T2605, T2634, and T2643 in mouse, respectively) are conserved (Fig. 1A) (20, 29, 30). To interrogate the impact of DNA-PKcs T2609 cluster phosphorylation, we replaced all five threonine residues in this region with alanines and named the allele -5A. Similar to the previously published 3A allele (which replaces the three conserved threonines)(33), the *DNA-PKcs^5A/5A^* mice were born alive with hyper-pigmented toes, but succumbed to bone marrow failure by 3 weeks with very few mature lymphocytes (35)(Fig. 1B, and Supplementary Fig. 1A and 1B). *Tp53*-deficiency, both heterozygous and homozygous, rescues the growth retardation, corrects the hyperpigmentation, and most importantly, restores peripheral splenic B cell frequency (Fig.1B, Supplementary Fig. 1A, and 1B), consistent with the dispensible role of T2609 phosphorylation in V(D)J recombination (35, 36).

### *DNA-PKcs^5A/5A^* B cells undergo slightly delayed, but efficient Class Switch Recombination

To understand the role of DNA-PKcs T2609 phosphorylation in CSR, we purified splenic B cells from *DNA-PKcs^5A/5A^Tp53^+/-(or -/-)^* and control *DNA-PKcs^+/+^* or *Tp53^+/-(or -/-)^* mice and cultured them with anti-CD40 and interleukin 4 (IL-4) that induce robust CSR to IgG1 and IgE. *Tp53*-deficiency does not affect CSR efficiency or CSR junctions (16, 17, 44, 45). About 20% *DNA-PKcs^+/+^* B cells were IgG1+ after 3 days of stimulation and ~37% IgG1+ by day 4 (Fig.1C and 1D). In comparison, *DNA-PKcs^5A/5A^Tp53^+/-(or -/-)^* B cells achieved ~15% IgG1^+^ B cells by day 3 and eventually 30-40% IgG1^+^ by day 4 of activation (Fig.1C and 1D). Successful CSR requires cell proliferation. Cell division analyses *via* the Cell Trace Violet (CTV) dye did not find proliferation defects in *DNA-PKcs^5A/5A^Tp53^+/-(or -/-)^* B cells (Supplementary Fig. 1C), consistent with the largely normal weight and peripheral lymphocyte numbers of *DNA-PKcs^5A/5A^Tp53^+/-(or -/-)^* mice (Supplementary Fig. 1A) (35). These data suggest that phosphorylation of DNA-PKcs at the T2609 cluster is not required for efficient CSR, despite a moderate delay in kinetics.

### T2609 cluster phosphorylation alters the distribution of the IgH junctions

Given *DNA-PKcs^-/-^* B cells undergo efficient CSR to IgG1 via the Alt-EJ mediated repair pathway, as detected by the HTGTS of Sμ-Sμ internal deletions and Sμ-Sγ1 CSR junctions (16), we decided to analyze the CSR junctions in *DNA-PKcs^5A/5A^p53^+/-(or -/-)^* B cells using the sensitive HTGTS. Briefly “translocation junctions” from a single bait site at the 5’Sμ to various targets were isolated after linear amplification and sequenced (Fig. 2A) (16, 41). Altogether, we analyzed 4545 junctions from B cells derived from 5 different *DNA-PKcs^5A/5A^p53^+/-(or -/-)^* mice and over 10,000 junctions total from control *DNA-PKcs^+/+^*, *Tp53^+/-^* and *DNA-PKcs^-/-^* mice. As shown in Figure 2B, nearly 90% of all junctions isolated from *DNA-PKcs^+/+^*, *Tp53^-/-^* or *Tp53^+/-^*, *DNA-PKcs^5A/5A^p53^+/-(or -/-)^* and *DNA-PKcs^-/-^* mature B cells are joined with a target (referred as “Prey”) within the IgH locus (defined as a 400kb region including all D, J elements and Constant regions). We further defined the orientation of Prey junctions as (+) if it reads from centromere to telomere after the junction, or as (–) if it reads from telomere to centromere. Since the IgH locus resides on the (-) strand of murine chromosome 12 and the bait at the 5’ Sμ reads from telomere to centromere, the normal internal deletion (Sμ-Sμ) and class switch recombination (Sμ-Sγ1, or Sμ-Sε) junctions are in the (-) orientation (Fig. 2A). Indeed, > 80% of all preys from *DNA-PKcs^+/+^* (WT) and *Tp53*-deficient B cells fall on the (-) strand of IgH (83% for WT and 82% for *Tp53*-deficient). Meanwhile, only 10% and 11% of all preys fall on the (+) strands of IgH in the *DNA-PKcs^+/+^* (WT) and *Tp53*-deficient cells (Fig. 2A). In contrast, only 75% of all preys from *DNA-PKcs^5A/5A^* B cells are on the IgH (-) strand and an increased 17% fall on the IgH (+) strand, similar to those in control *DNA-PKcs^-/-^* B cells (68% IgH+ and 22% IgH-) (Fig. 2B and 2C) (16). To understand the significance of the relative increases of the IgH+ preys among all IgH preys, we examined the orientation of the inter-chromosomal preys outside of the IgH locus (almost all to another chromosome beyond chromosome 12). Non-IgH (nIgH) preys fall equally on (+) and (-) strands in all genotypes tested (Fig. 2B), suggesting that the negative strand bias among the IgH preys reflects a strong preference for intra-chromosomal deletion joining. Thus, we hypothesized that the relative increase of IgH preys falling on the (+) strand in *DNA-PKcs^5A/5A^* and *DNA-PKcs^-/-^* B cells reflect either increased inversional events (Fig 2C) (as illustrated with blue arrows in Fig. 2A) or increased inter-sister chromatid or inter-chromosomal translocations (between IgH of two homologous chromosomes 12). A similar phenotype has been noted for B cells lacking cNHEJ factors (e.g., *Xrcc4^-/-^*) or DNA damage response factors (*e.g., Atm^-/-^* and *Tpr53BP1^-/-^*) (16, 46, 47).

**Figure 2:**
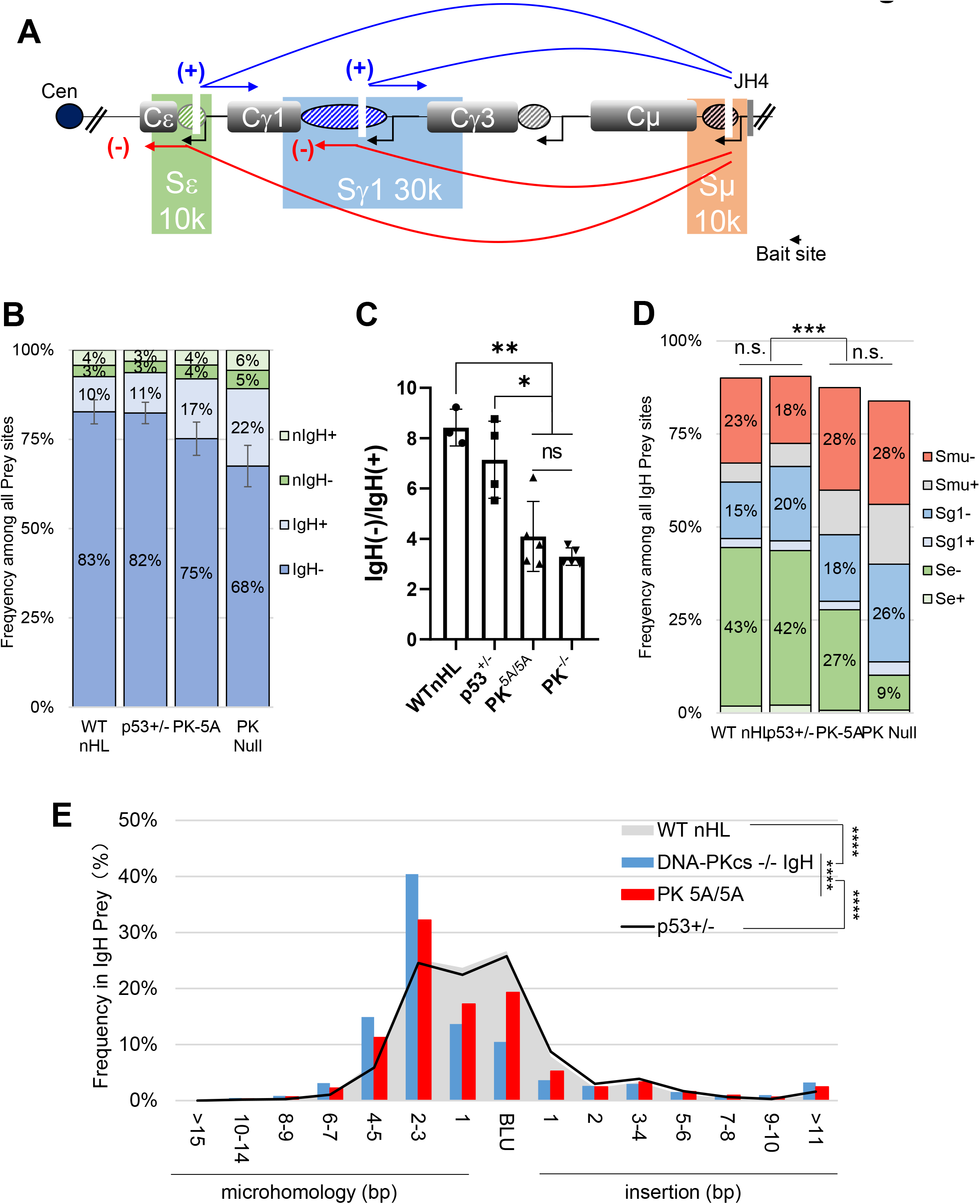
Class switch recombination in *DNA-PKcs^5A/5A^* cells have increased MH. (A) Diagram of the murine IgH locus with the location of Sμ (beige), Sγ1 (blue) and Sε (green). The bait site is marked on the lower right of the diagram. The red (deletional, (-) strand) and blue (inversional, (+) strand) arrows indicate the orientation of potential junctions. (B) Relative frequency of HTGTS junctions in the (-) and (+) strand orientation, and in the IgH and nonIgH regions. We defined the junction as (+) orientation if it reads from the centromere to the telomere, and as (-) if it reads from the telomere to the centromere. The IgH region is defined as 400kb including part of V, all D, all J segments and the entire switch region. The error bars represent standard deviations. (C) Ratio between total number of junctions mapped to (-) strand of IgH and (+) strand of IgH, The p <0.005 **, p<0.05 *, n.s. (Not significant) for unpaired student’s t-test between the IgH(-) prey%. (D) Frequency of HTGTS junctions in the (-) and (+) strand orientation, and in the IgH regions, subdivided into Sμ, Sγ1, and Sε. The data represent the sum of data from 5 *DNA-PKcs^5A/5A^* mice, 3 *DNA-PKcs^+/+^* mice (WT), 4 *DNA-PKcs^-/-^* and a total of 4 (2 each) *Tp53^-/-^* or *Tp53^-/-^* mice. The p <0.001 ***, n.s. (Not significant) for unpaired student’s t-test. (E) Frequency of junctions with microhomologies (MH), Blunt (BLU), and insertions (INS). The p <0.0001 ****, n.s. (Not significant) for Kolmogorov-Smirnov test.

The cytokine combination anti-CD40 and IL-4 induce CSR to IgG1 and IgE. Accordingly, ~42% of all junctions from *DNA-PKcs^+/+^*, and *Tp53^-/-^* or *Tp53^+/-^* mice were located in Sε (-) (potentially IgE switching), 15~20% in Sγ1(-) (potential IgG1 switching) and 18-23% in Sμ (-) (internal deletion) (Fig. 2D). IgH junctions from both *DNA-PKcs^-/-^* and *DNA-PKcs^5A/5A^* B cells have a greatly increased Sμ (-) portion (28%), and a corresponding reduction in Sε (-) preys (9% and 27%, respectively) (Fig.2D). The overall proportion changes are more dramatic in those from *DNA-PKcs^-/-^* B cells than those from *DNA-PKcs^5A/5A^* B cells (16). Together these data suggest that despite normal CSR efficiency, the junctions from *DNA-PKcs^5A/5A^* B cells contain increased translocations (to the (+) strand) and preferential loss of Sμ-Sε junctions – both features of cNHEJ deficiency, similar to those from *DNA-PKcs^-/-^* cells. Correspondingly, Florenscence In situ Hybridization with a telomere probe to increase the sensitivity to IgH locus breaks near the telomere described before (48) also revealed increased genomic instability, including both chromosome breaks at both sister chromatids, and chromatid breaks in activated *DNA-PKcs^5A/5A^* B cells (Fig.S2A, S2B, and S2C)

### Loss of T2609 cluster phosphorylation increase MH usage in CSR junctions

In the absence of cNHEJ, including the loss of DNA-PKcs, significant CSR is achieved via the Alt-EJ pathway that preferentially uses MHs at the junction (16–18). In a recent analysis, we identified two features associated with Alt-EJ dependent CSR using HTGTS (16) - increased usage of MH at the junctions and evidence of extensive resection in switch regions. Indeed, junctions from *DNA-PKcs^-/-^* B cells with only moderate CSR defects contain both features (14, 15). Among CSR junctions (including all IgH preys) recovered from *DNA-PKcs^+/+^* or *Tp53*-deficient (alone) cells, ~25% are blunt, 25% have 1nt MH and another 25% have 2-3nt MH. In contrast, less than 15% of junctions from *DNA-PKcs^-/-^* and 20% from *DNA-PKcs^5A/5A^* cells were blunt and only another 15% have 1nt MH (Fig. 2E). A prominent 40% of junctions from *DNA-PKcs^-/-^* B cells and 32% of the IgH junctions from *DNA-PKcs^5A/5A^* B cells have 2-3nt MH. Indeed, cumulatively, nearly 20% of IgH junctions from *DNA-PKcs^-/-^* and *DNA-PKcs^5A/5A^* B cells have >4 nt MH (in contrast to ~10% in *DNA-PKcs^+/+^* or *Tp53*-deficient alone) (Fig. 2E). The overall frequency of insertions within IgH junctions did not differ between genotypes (Fig.2E). Together these findings indicate that loss of the T2609 phosphorylation of DNA-PKcs unleashed robust MH-mediated CSR that compensates for the moderate loss of cNHEJ.

### Loss of T2609 cluster phosphorylation increases end-resection

Next, we mapped the junctions within and around each specific switch region. The (+) and (-) oriented junctions were plotted above (blue) and below (red) the centerline, respectively (Fig, 2A, 3, 4, and 5). Since the location of the bait is fixed within the 5’ Sμ (telomeric), the majority of the Sμ joins from *DNA-PKcs^+/+^* and *Tp53*-deficient B cells are located near the bait site only ~10% beyond the core Sμ region (Fig.3). In contrast, over 15% of Sμ junctions from *DNA-PKcs^5A/5A^* B cells fall beyond the core Sμ region, a similar trend to what has previously been described in *DNA-PKcs^-/-^* cells (Fig. 3). *DNA-PKcs^5A/5A^* junctions show a similarly wider distribution within Sγ1, with 1.66% falling downstream of Sγ1 (versus 0.14-0.54% in the controls) (Fig. 4), and within Sε (9.13% in *DNA-PKcs^5A/5A^* junctions versus ~6% in the controls) (Fig. 5). Altogether, the extensive use of distal switch regions suggests end-resection, another feature of Alt-EJ, and cNHEJ deficiency in *DNA-PKcs^5A/5A^* B cells.

**Figure 3:**
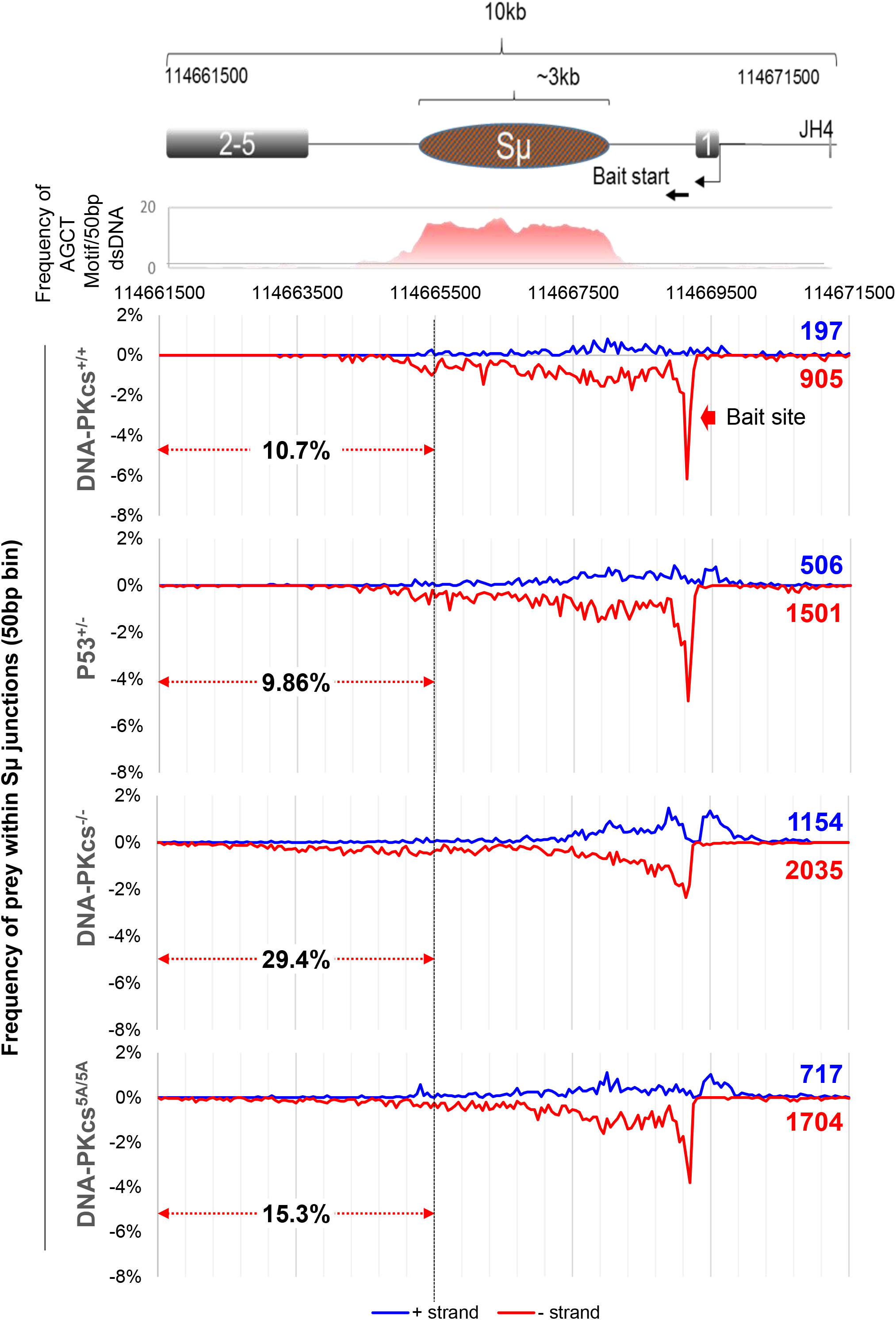
Distribution of CSR junctions within 10kb of the Sμ locus. The schematic of the region of interest is drawn at the top. For each genotype, the number of junctions is indicated for the (+) strand (blue) and (-) strand (red). The (-) orientation junctions are shown as negative percentages. Percent of junctional usage from the end of Sμ to the end of the 10kb window (between chr12:114661500 – 114665500) is indicated with red arrows. Junctions in this range are considered resections. Bin=50bp. Only the percentage of negative strand junctions was used to calculate the resection %. The data represent the pool of 5 *DNA-PKcs^5A/5A^* mice, 3 *DNA-PKcs^+/+^* mice (WT), 4 *DNA-PKcs^-/-^* mice, and a total of 4 (2 each) *Tp53^-/-^* or *Tp53^+/-^* mice.

**Figure 4:**
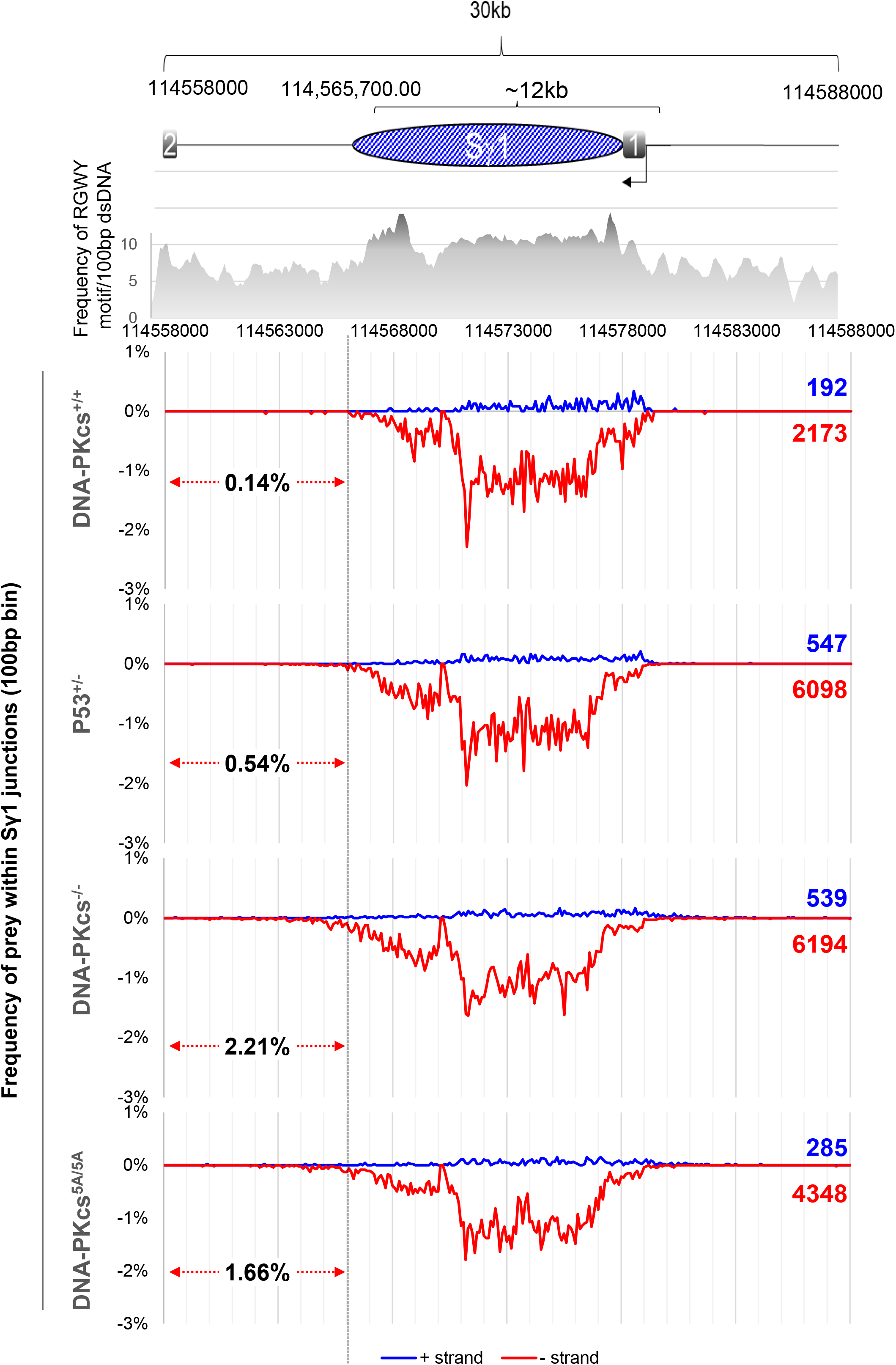
Distribution of CSR junctions within 30kb of the Sγ1 locus. The schematic of the region of interest is drawn at the top. For each genotype, the number of junctions is indicated for the (+) orientation (blue) and (-) orientation (red). The (-) orientation junctions are shown as negative percentages. Percent of junctional usage from the end of Sγ1 to the end of the 30kb window (between chr12: 114558000 – 11456600) is indicated with red arrows (bin=100bp). Junctions in this range are considered resections. Only the percentage of negative strand junctions was used to calculate the resection %. The data represent the pool of 5 *DNA-PKcs^5A/5A^* mice, 3 *DNA-PKcs^+/+^* mice (WT), 4 *DNA-PKcs^-/-^* mice, and a total of 4 (2 each) *Tp53^-/-^* or *Tp53^+/-^* mice.

**Figure 5:**
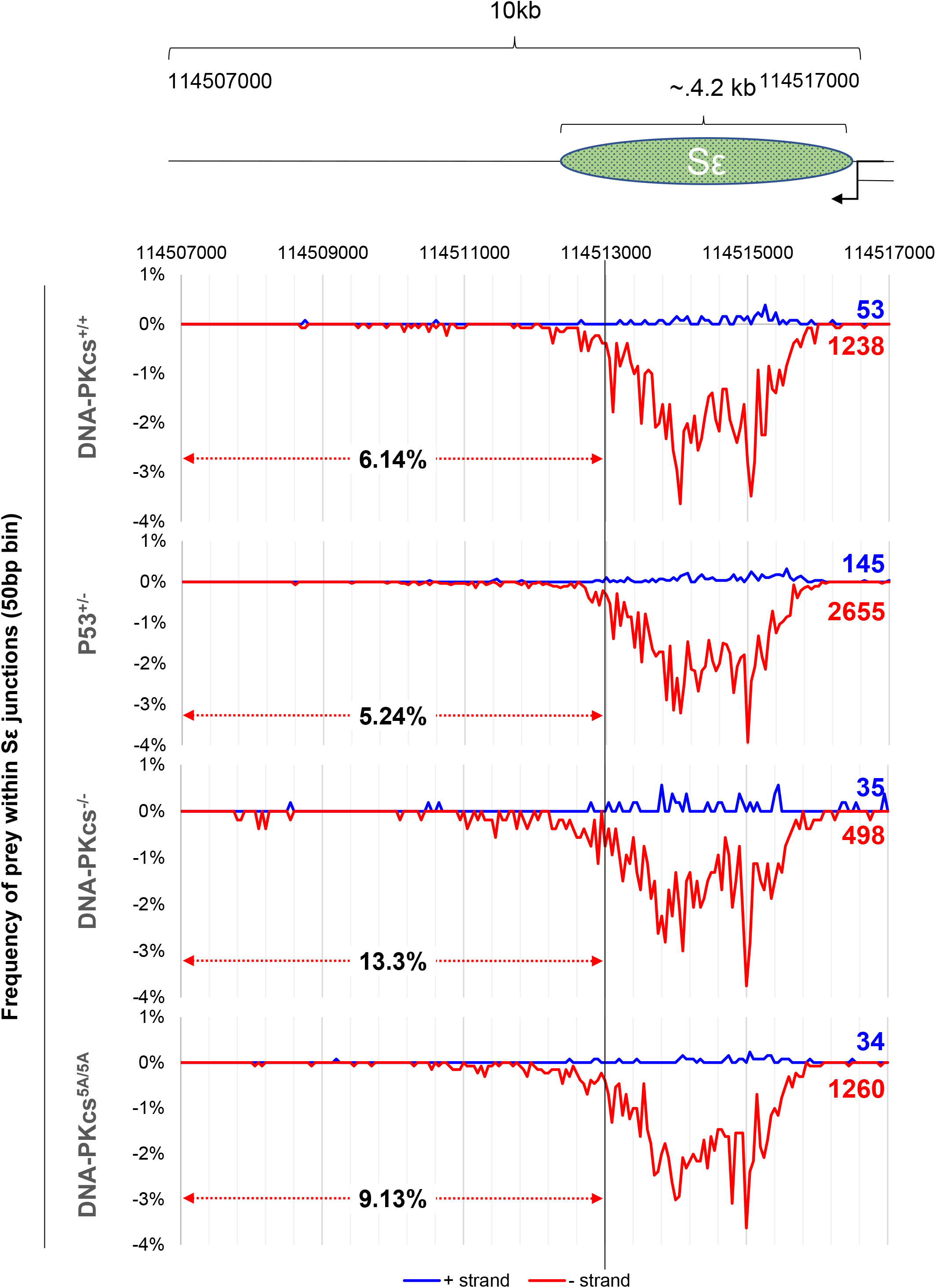
Distribution of CSR junctions within 10kb of the Sε locus. The schematic of the region of interest is drawn at the top. For each genotype, the number of junctions is indicated for the (+) strand (blue) and (-) strand (red). The (-) orientation junctions are shown as negative percentages. Percent of junctional usage from the end of Sε to the end of the 10kb window (between chr12:11450700 – 114513000) is indicated with red arrows (50bp bin). Junctions in this range are considered resections. Only the percentage of negative strand junctions was used to calculate the resection %. The data represent the pool of 5 *DNA-PKcs^5A/5A^* mice, 3 *DNA-PKcs^+/+^* mice (WT), 4 *DNA-PKcs^-/-^* mice, and a total of 4 (2 each) *Tp53^-/-^* or *Tp53^+/-^* mice.

### Loss of T2609 phosphorylation reduced G-mutations in the 5’Sμ bait region of IgH

Finally, we analyzed mutations in the 5’Sμ region among the junctions (Fig. 6). To compare with prior studies using Sanger sequencing, we present the data based on the top strand relative to the Sμ germline transcription (Fig. 6A). The estimated mutation rate for the MiSeq platform is 0.1% (10 mut/10,000nt). The overall mutation rate (mut/10,000nt) and frequency of reads with mutations were both higher in IgH junctions derived from *DNA-PKcs^+/+^* and *Tp53^-/-^* cells (~27 mut/10,000nt) than those from *DNA-PKcs^5A/5A^* cells (~16 mut/10,000nt) that were also Tp53 deficient (~16 mut/10,000nt) (Fig. 6D). Junctions from *DNA-PKcs^-/-^* cells also had a slightly lower mutation rate than *DNA-PKcs^+/+^* and *Tp53^-/-^* cells (Fig. 6D). Despite a similar distribution of nucleotide types (Fig. 6B), G was most frequently mutated in the 5’Sμ region of *DNA-PKcs^+/+^* and *Tp53^-/-^* B cells (47 mut/10,000 G). Lower G mutation rates in *DNA-PKcs^-/-^* cells (20 mut/10,0000 G) and *DNA-PKcs^5A/5A^* cells (25mut/10,000 G) account for a lower overall mutation rate in those cells (Fig. 6F). The pattern of G mutations (relative frequency to A, T, C) did not change, suggesting the reduction is likely due to AID targeting, rather than differences in secondary processing by uracil-DNA glycosylase (UNG), Mismatch repair (MMR) or other pathways (Fig. 6C). Moreover, there is also an overall reduction in the frequency of reads with mutations in *DNA-PKcs^5A/5A^* cells (Fig. 6E). Altogether, we note a reduction of G nucleotide mutations in the 5’Sμ region in *DNA-PKcs^5A/5A^* and DNA-PKcs^-/-^ cells. Similar changes have been noted in cNHEJ deficient, *Xrcc4^-/-^* and *DNA-PKcs^KD/KD^* B cells.

**Figure 6:**
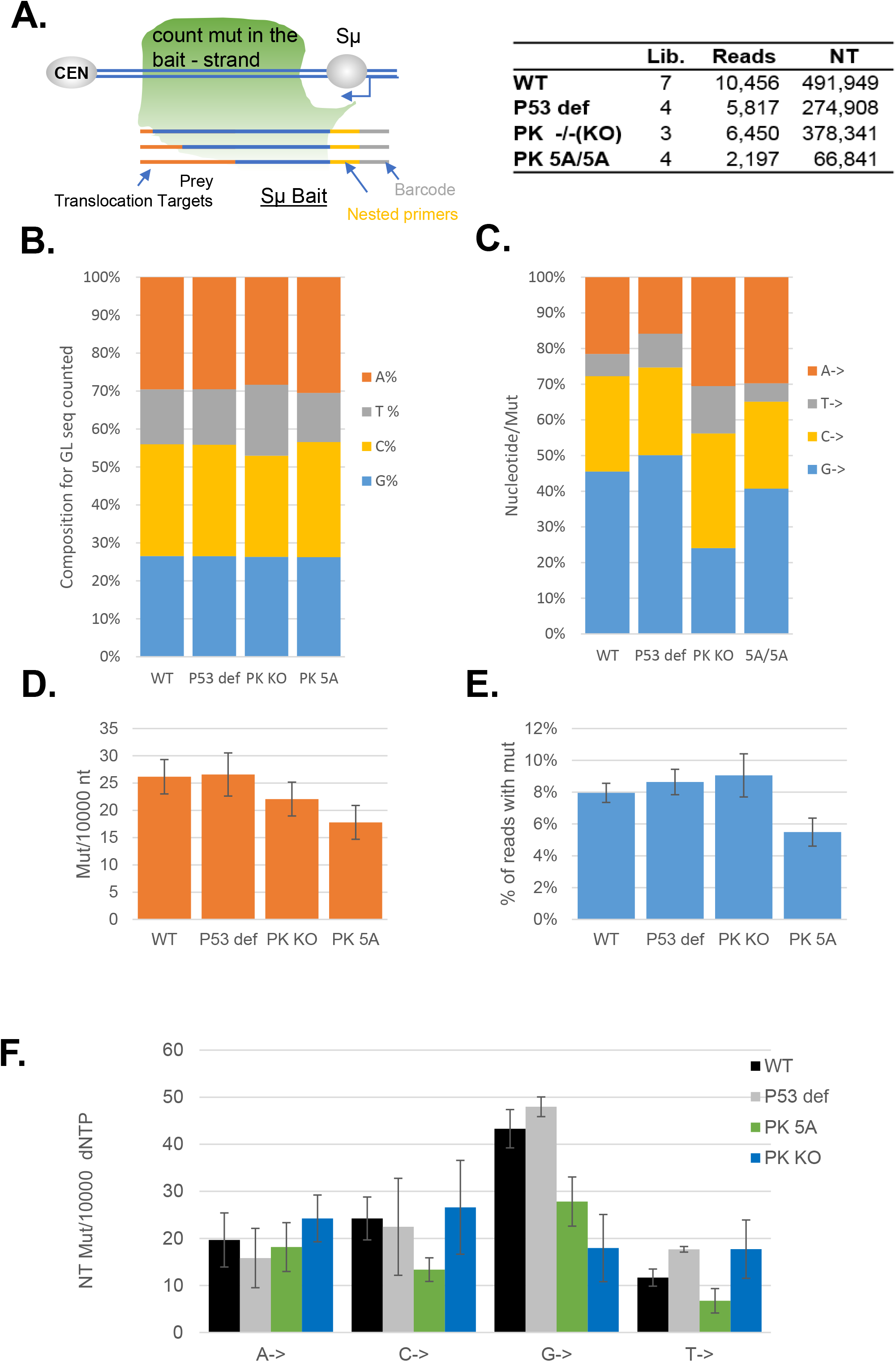
*DNA-PKcs^5A/5A^* HTGTS CSR junctions do not have a strong mutational pattern in the IgH region. (A) Diagram of the bait site and nucleotide counting scheme for mutational analysis (left). For mutation analyses, we counted the mutations after the nested PCR primer until the end of the bait sequence. There is a clear difference between mutation frequency in IgH junctions vs non-IgH junctions even if only the bait sequence is compared, so only IgH junctions (both + and – orientation) were counted. The number of independent libraries, reads, and nucleotides analyzed for mutational distribution is shown on the right. (B) Distribution of nucleotide (A, T, C, or, G) frequency in the bait portion used to calculate mutation frequency. (C) Frequency of mutations of each nucleotide (A, T, C, or G) in all mutations obtained. (D) The overall frequency of mutations per 10,000 nucleotides (nt). (E) Frequency of all IgH junctions (reads) containing >0 mutations. (F) Mutations in a particular nucleotide per 10,000 of that nucleotide type. The error bars represent standard errors. The data represent the collection from 4 *DNA-PKcs^5A/5A^* mice, 4 *DNA-PKcs^+/+^* mice (WT), 4 *DNA-PKcs^-/-^* and 2 *Tp53^+/-^* mice.

## Discussion

DNA-PKcs deficiency has been linked to reduced CSR efficiency and a preference for MH in patients and mouse models(16, 19). The T2609 cluster phosphorylation is one of the most prominent post-translational modification of DNA-PKcs and has been associated with increased radiation sensitivity (22, 28–31), which its role in CSR remains elusive. In this context, the T2609 phosphorylation has been implicated in telomere biology (34) and ribosomal biogenesis (35) which both could indirectly contribute to radiation sensitivity. To understand the role of T2609 phosphorylation in CSR and DSB repair specifically, we evaluated the role of T2609 phosphorylation in physiological DSB repair during CSR using germline mouse models. As in the case of V(D)J recombination (33, 35, 36), T2609A mutation also does not affect CSR efficiency. But CSR junctions from *DNA-PKcs^5A/5A^* B cells revealed a preferential loss of Sμ-Sε junctions, increased MH usage, and evidence of extensive end resection and increased inter-chromosomal translocations ((+) strand preys)–all consistent with cNHEJ deficiency and compensatory Alt-EJ-mediated CSR. Together these findings support the role of DNA-PKcs T2609 phosphorylation in promoting cNHEJ-mediated CSR. In the absence of T2609 phosphorylation, the increased dependence on the Alt-EJ pathway, together with the extensive length of the Sγ1 region (12.5kb in 129SV strain), the dense RGYW/AGCT sequence motifs (Fig. 4), and the sequence-independent 3D interactions (synapses) between Sμ-Sγ1 (49–52), lead to robust Alt-EJ-mediated CSR to IgG1 in *DNA-PKcs^5A/5A^* B cells. Thus, our analyses identify a novel role of DNA-PKcs T2609 phosphorylation in repair pathway choice during CSR.

The estimated error rate of the Illumina Miseq platform is 0.1%, which is lower than the estimated and reported somatic hypermutation (SHM) rate (53) and therefore allows for analyses of SHM. As previously reported (54, 55), G mutations are two times more frequent than C mutations on the top strand of the 5’Sμ region (the non-template strand relative to Sμ germline transcription). Interestingly, the T2609A mutation attenuated this preference of G mutations, while the pattern of G mutations (relative to A, C, T) is unaffected, suggesting that the T2609A mutation and its associated repair defects might compromise AID targeting to 5’Sμ without affecting U:G mismatch processing. Consistent with this model, the frequency of reads with mutations also decreased in *DNA-PKcs^5A/5A^* B cells. The bias of APOBEC family enzymes to deaminate the lagging strand during DNA replication (53, 56) and the relative abundance of sense vs anti-sense transcripts (57) could all contribute to this strand bias.

The CSR junctions from *DNA-PKcs^5A/5A^* and *DNA-PKcs^-/-^* cells share many features, including the end-resection and increased MH. T2609 phosphorylation is also dispensible for hairpin opening and V(D)J recombination(33, 35, 36, 40), suggesting a model in which the T2609 cluster phosphorylation promotes end-ligation, but is dispensable for end-processing and Artemis recruitment. The increased use of MHs and end-resection in *DNA-PKcs^5A/5A^* cells are also much less prominent than those from *DNA-PKcs^KD/KD^* cells (16, 40). Together with the normal CSR in *DNA-PKcs^PQR/PQR^* B cells with alanine substitutions at the S2056 cluster(32), these results suggest that lack of DNA-PKcs phosphorylation alone is not sufficient to explain the strong cNHEJ-defects in *DNA-PKcs^KD/KD^* cells. The possibilities include other phosphorylation sites (23) or phosphorylation-independent effects of catalysis itself on the structure or organization of DNA-PKcs protein. Recent structural analyses of DNA-PK holoenzymes have provided evidence for allosteric changes upon DNA-PK activation (58–60), which might link the catalysis directly with DNA-PKcs turnover and its end-protection function.

In summary, our findings unequivocally demonstrate an important distinction between auto-phosphorylation and catalysis on DNA-PKcs, indicating that auto-phosphorylation at T2609, is largely dispensable for lymphocyte development. Despite the robust CSR efficiency, the CSR junctions recovered from *DNA-PKcs^5A/5A^* cells reveal a strong bias toward Alt-EJ, suggesting a role of T2609 in promoting cNHEJ at the price of Alt-EJ during CSR.

## Acknowledgment

We thank Dr. Wenxia Jiang for her input on the DNA-PKcs KD mouse model and Ms. Verna Estes for assistance with the colony management. We thank Dr. Chyuan-Sheng (Victor) Lin for technical assistance and advice on the generation of *DNA-PKcs^5A/5A^* mouse models. We thank other members of the Zha lab for helpful discussions and technical advice. We apologize to colleagues whose work could not be cited due to space limitations and was covered by reviews instead.

## Footnotes

### 1) Grant Support

This work is in part supported by NIH/NCI 5R01CA158073, 5R01CA184187, and R01CA226852. SZ is the recipient of the Leukemia Lymphoma Society Scholar Award. JC was supported by NIH/NCI F31CA183504-01A1 and CTSA/NIH TL1 TR000082. This research was funded in part through the NIH/NCI Cancer Center Support Grant P30CA013696 to the Herbert Irving Comprehensive Cancer Center (HICCC) of Columbia University.

### 2) Author Contributions

JC, XW, SZ designed experiments. SZ wrote the paper with help from XW and JC. JC and XW analyzed the DNA-PKcs 5A mice and cells for lymphocyte development and function. JC generated the DNA-PKcs 5A mice. XW, JC, and BL established the HTGTS assay and performed the HTGTS analyses of the CSR junctions for this study. VE helped with the breeding and animal care.

## 3) Abbreviations

Alt-EJ: alternative end-joining
cNHEJ: classical non-homologous end-joining
CSR: class switch recombination
DNA-PK: DNA-dependent protein kinase
DSB: double-strand break
HTGTS: High throughput genomic translocation sequencing
MH: micro-homology
SHM: somatic hypermutation

---

**SupFigure 1.**
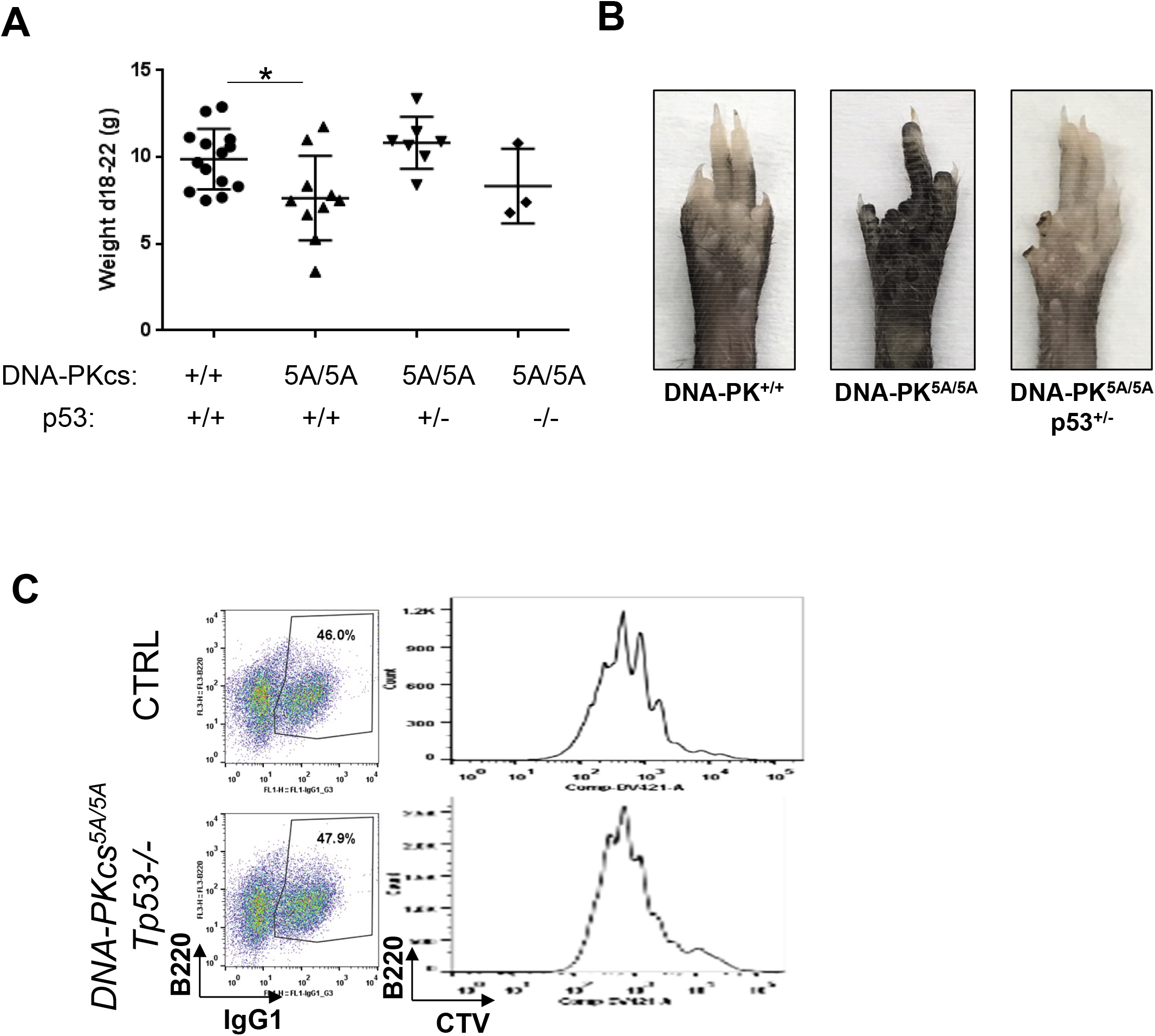

**SupFigure 2.**
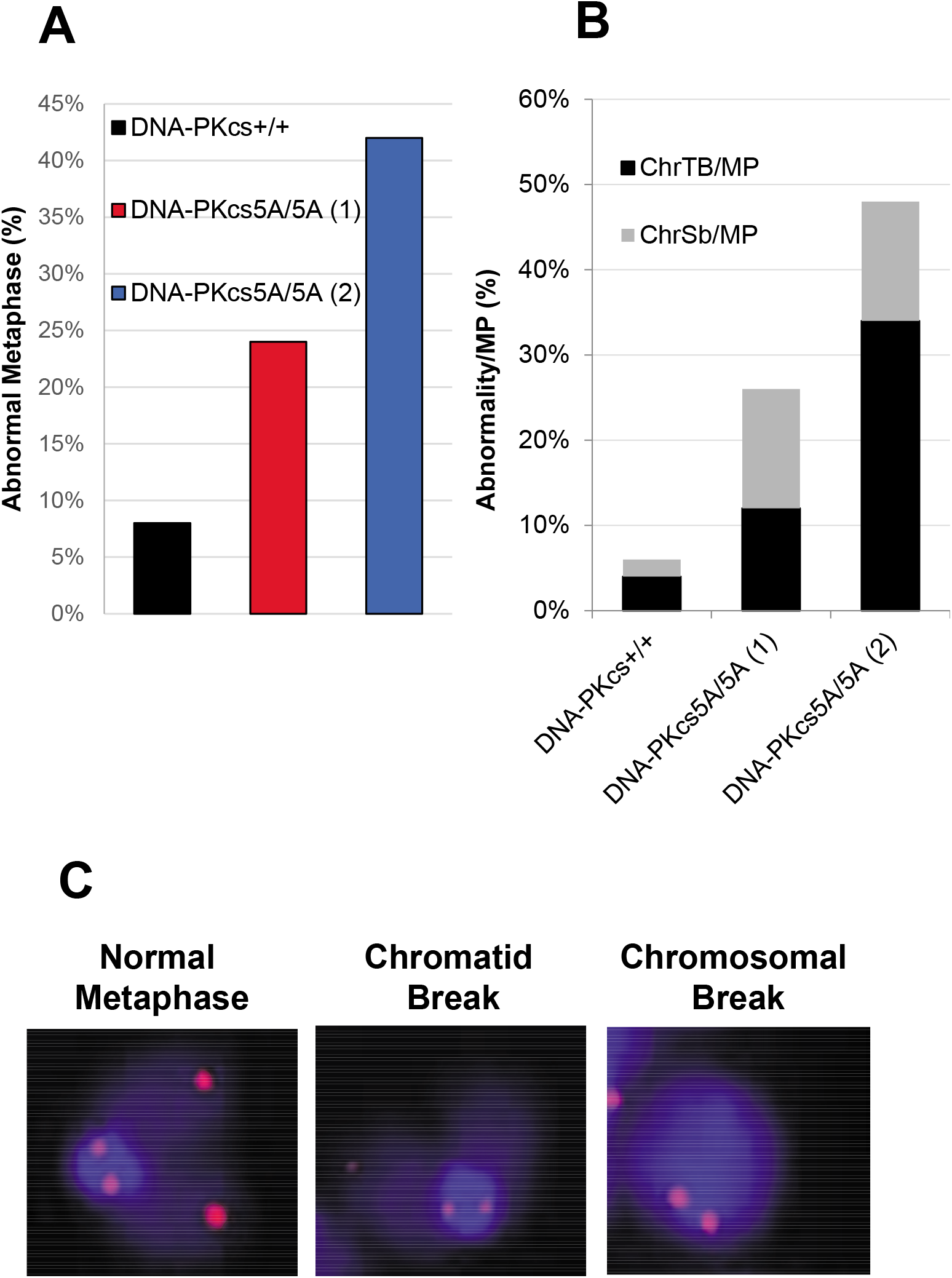

## Notes

### Competing Interest Statement

The authors have declared no competing interest.

